# A novel adjuvant formulation induces robust Th1/Th17 memory and mucosal recall responses in Non-Human Primates

**DOI:** 10.1101/2023.02.23.529651

**Authors:** Joshua S Woodworth, Vanessa Contreras, Dennis Christensen, Thibaut Naninck, Nidhal Kahlaoui, Anne-Sophie Gallouët, Sébastien Langlois, Emma Burban, Candie Joly, Wesley Gros, Nathalie Dereuddre-Bosquet, Julie Morin, Ming Liu Olsen, Ida Rosenkrands, Ann-Kathrin Stein, Grith Krøyer Wood, Frank Follmann, Thomas Lindenstrøm, Roger LeGrand, Gabriel Kristian Pedersen, Rasmus Mortensen

## Abstract

After clean drinking water, vaccination is the most impactful global health intervention. However, development of new vaccines against difficult-to-target diseases is hampered by the lack of diverse adjuvants for human use. Of particular interest, none of the currently available adjuvants induce Th17 cells. Here, we develop and test an improved liposomal adjuvant, termed CAF®10b, that incorporates a TLR-9 agonist. In a head-to-head study in non-human primates (NHPs), immunization with antigen adjuvanted with CAF®10b induced significantly increased antibody and cellular immune responses compared to previous CAF® adjuvants, already in clinical trials. This was not seen in the mouse model, demonstrating that adjuvant effects can be highly species specific. Importantly, intramuscular immunization of NHPs with CAF®10b induced robust Th17 responses that were observed in circulation half a year after vaccination. Furthermore, subsequent instillation of unadjuvanted antigen into the skin and lungs of these memory animals led to significant recall responses including transient local lung inflammation observed by Positron Emission Tomography-Computed Tomography (PET-CT), elevated antibody titers, and expanded systemic and local Th1 and Th17 responses, including >20% antigen-specific T cells in the bronchoalveolar lavage. Overall, CAF®10b demonstrated an adjuvant able to drive true memory antibody, Th1 and Th17 vaccine-responses across rodent and primate species, supporting its translational potential.

## INTRODUCTION

Adjuvants are immunostimulating components that are included in vaccines to enhance the magnitude, phenotype or longevity of the resulting immune responses. With scientific advancements, the list of licensed vaccines using adjuvants has expanded during the last decade and now includes subunit vaccines against human papillomavirus, hepatitis B, pandemic influenza, malaria, shingles disease and SARS-CoV-2. However, since the introduction of alum in the 1920s, only a handful of new adjuvants have become available for human use (*1*). This significantly hampers the development of novel vaccines and there is a recognized need for diversification in the portfolio of available adjuvants/vaccine platforms (*2, 3*).

In the ideal scenario, different adjuvants could be used to tailor the immune responses for specific needs, but the mechanisms by which many of the existing adjuvants work and the functional responses they induce are only partially understood (*1*). For instance, few of the available adjuvants were initially designed to induce robust T cell responses, and it has proven challenging to specifically direct cellular immune functionalities. Of particular interest, the existing human adjuvants have not been shown to induce a detectable Th17 response, despite the growing evidence of a protective role of Th17 cells against both viral and bacterial infections, such as influenza, *Bordetella pertussis* and *Mycobacterium* tuberculosis (Mtb) (*4–12*). Therefore, together with increasing the magnitude of cellular immune responses, induction of Th17 cells is a prime goal for vaccine optimization. Another limitation of some of the most successful current adjuvants is that active components are derived from natural sources. This can create bottlenecks for product availability, as seen with QS21 that is dependent on soap bark trees (*13*) and squalene that is sourced from shark liver (*14*). Alternatively, with a greater understanding of immune signaling pathways and advancements in synthetic chemistry, there is a possibility to design highly specialized adjuvants that can be manufactured from widely available constituents.

The cationic adjuvant formulation (CAF®) family consists of well-defined stable liposomes that facilitate co-delivery of antigen and immunostimulator to target cells and slow antigen release by forming depots at the injection site (*15*). They consist of all-synthetic components, making the CAF® platform an ideal starting point for developing the next generation of safe and immunogenic adjuvants. Furthermore, several CAF® adjuvants are able to stimulate Th17 responses after subcutaneous immunization in mice (*15, 16*), a feature not observed for licensed adjuvants. The first member of the family, CAF®01, consisting of dimethyl-dioctadecylammonium (DDA) and the Mincle agonist Trehalose-6,6-dibehenate (TDB), has completed 5 clinical trials and has been shown to induce 5-6 fold higher IgG titers than alum and stable cell-mediated immune responses for more than 150 weeks in humans (*17, 18*). However, for some disease targets, a stronger cell-mediated immune (CMI) response is desirable, and encouraged by the clinical success of the highly immunogenic AS01® adjuvant in shingles, malaria and TB vaccines (*19, 20*), the goal of this study was to develop and test a next generation CAF® adjuvant displaying increased long-term memory responses, while also introducing vaccine-priming of Th17 cells. Such an adjuvant would have a broad range of applications, including vaccines for respiratory infections, where Th17 cells are described to form resident memory T cells and accelerate local immune responses (*21, 22*). It would therefore also be the ideal candidate to be co-developed with the recently introduced tuberculosis vaccine antigen, H107, from our group (*23*).

Here we introduce a new potent CAF® adjuvant formulation, CAF®10b, that incorporates a TLR9 agonist and is shown to induce robust Th17 responses in both mice and non-human primates (NHPs). CAF®10b is compared head-to-head in mice and NHPs against three other adjuvants; *i)* CAF®01, included as a benchmark adjuvant with clinical data and comparative results to other human-approved adjuvants (*17, 18, 24, 25*), *ii)* CAF®09b, which incorporates the TLR3 agonist poly(I:C) and is in clinical trials for cancer immunotherapy (*26*) and *iii)* a higher-dose formulation of CAF®09b (CAF®09^hi^). Our study shows that CAF®01 and CAF®10b gave the highest cellular immune responses in mice, including significant Th1 and Th17 responses. In contrast, CAF®10b provided the highest Th1 and Th17 memory responses in NHPs. Importantly, the CAF®09^hi^ formulation gave the highest peak cellular (Th1) response, but a lower detectable memory response than CAF®10b, demonstrating that peak cellular immune responses do not necessarily reflect memory responses. At six months after vaccination, we used *in vivo* antigen recall to study functional memory, and here the CAF®10b-induced T cells showed potent systemic expansion and ability to migrate into local tissues, including the lung mucosa. These immunogenicity data for CAF®10b are highly encouraging and demonstrate the potential to fill an important gap in modern vaccine development.

## RESULTS

### Incorporation of CpG2006 into DDA/MMG makes stable liposomes that induce Th1/Th17 responses in mice

As a starting point for our efforts to develop a new CAF® formulation with increased CMI responses, we leveraged previous studies demonstrating a beneficial effect of incorporating CpG1826 in liposomes composed of dimethyldioctadecylammonium/monomycoloyl glycerol (DDA/MMG) in mice (*27*). As CpG1826 is a weak ligand for human TLR9, we explored the use of CpG2006, which has already been validated in human clinical trials (*28, 29*). Using HEK-293 cells expressing the human TLR9 gene and an NF-κB driven SEAP reporter, we confirmed that CpG2006 stimulated human TLR9 while CpG1826 did not (Fig. 1A). To our surprise, we also found that CpG2006 stimulated HEK-293 cells expressing murine TLR9.

**Figure 1.**
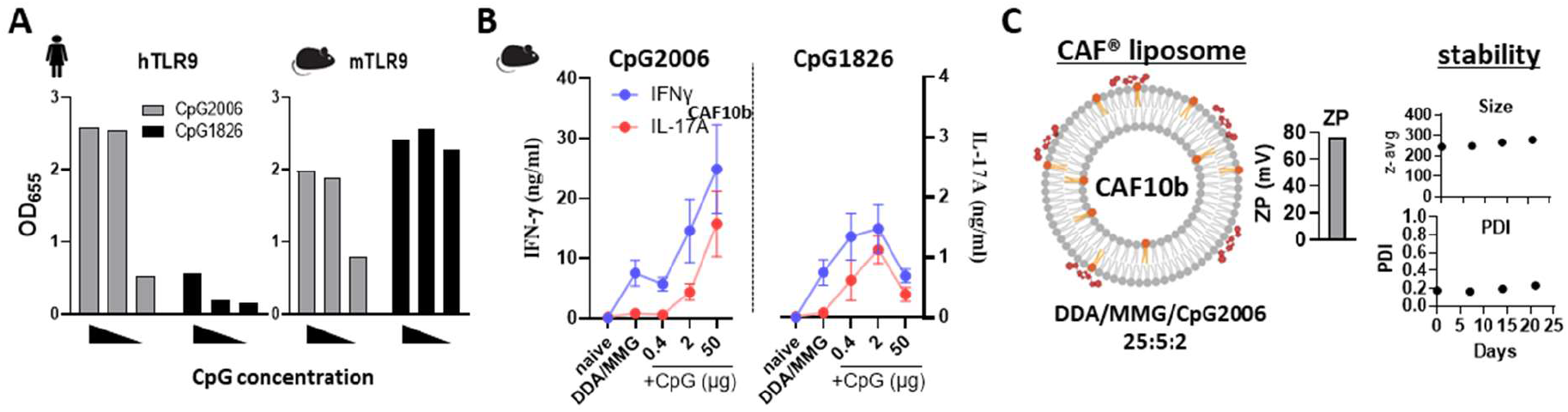
Incorporation of CpG2006 in CAF® generates stable liposomes that induce Th1/Th17 responses in mice. **(A)** HEK-Blue reporter cells expressing human TLR9 (left) or murine TLR9 (right) were stimulated in vitro with decreasing concentrations (100, 10, 1 μg/ml) of CpG2006 (grey bars) or CpG1826 (black bars) for 17 hrs and supernatants assayed by SEAP for HEK-Blue detection QUANTI-Blue™ (shown as OD_655_). The experiment was performed twice. **(B)** DDA/MMG liposomes were combined with various amounts of CpG2006 (left) or CpG1826 (right), adsorbed with protein antigen, and used to immunize CB6F1 mice 2x SC. Two weeks later, PBMCs stimulated *ex vivo* with protein antigen were assessed by ELISA for secretion of IFNγ (blue) and IL-17A (red). Symbols indicate mean±SD. **(C)** The depicted CAF®10b adjuvant was assessed for zeta potential (ZP) (left), particle size (top right) and polydispersity index (bottom right) over 25 days.

Given the ability of CpG2006 to activate murine TLR9, we next incorporated CpG2006 into CAF® liposomes and studied responses in mice. We first confirmed that incorporation of CpG2006 into the DDA/MMG liposomes significantly reduced systemic inflammation associated with administering free CpG2006 (Fig. S1A). We next studied the immunogenicity after vaccination. CB6F1 mice were subcutaneously (SC) immunized two times, three weeks apart, with antigen adsorbed to DDA/MMG liposomes with increasing CpG concentrations. Immune responses were analyzed two weeks after the last immunization by measuring IFNγ and IL-17 release from antigen-stimulated splenocytes. We observed that incorporation of CpG2006 increased both Th1 and Th17 responses (Fig. 1B, left), whereas CpG alone did not induce Th17 responses (Fig. S1B). There was also a clear dose dependency, with the highest dose of CpG (50μg) giving the highest responses (Fig. 1B, left). In contrast, with CpG1826 we observed the highest responses for the ‘intermediate’ dose of 2μg with lower responses for the 50μg dose, suggesting that overdosing of CpG is possible in target species where the TLR9 recognition is strong and is consistent with the tolerogenic effects of high dose CpG *in vitro* (*30*). We therefore aimed for an intermediate immunogenic dose of CpG2006 for the final formulation, to be used in NHPs/humans, and observed that a 25:5:2 DDA/MMG/CpG2006 ratio resulted in highly stable liposomes (denoted CAF®10b). Similar to existing CAF® adjuvants, the CAF®10b liposomes had an average size of 250 - 300 nm and a net positive charge (Fig. 1C, right). Importantly, long-term stability studies at 2-8°C demonstrated a minimal increase in liposome size and no decline in CpG, DDA, or MMG concentrations over a nine months period (Fig S2 A-E).

In summary, formulation of CpG2006 with DDA/MMG increases CAF®-induced Th1 and Th17 responses in mice.

### CAF®10b induces responses similar to CAF®01 in mice and protects against Mtb challenge

Having developed stable CAF®10b liposomes, we next utilized the recently developed TB vaccine antigen H107 (*23*) to make head-to-head comparisons against existing CAF® adjuvants in the mouse model (Fig. 2A). CAF®01 has previously been shown to induce antibody, Th1, and Th17 responses in mice (*24*) and was chosen for direct comparison to CAF®10b. In addition, two formulations of CAF®09, DDA/MMG liposomes with ‘low’ and ‘high’ doses of MMG and the TLR3 agonist Poly(I:C), were also included to investigate the relative adjuvant effects of TLR3 agonist incorporation into DDA/MMG liposomes for comparison (*31*). The adjuvants CAF®01 and CAF09^lcι^ (CAF09b) are currently in clinical development (*15, 18, 26*). All CAF® formulations induced H107-specific antibody responses (Fig. S3). Serum anti-H107 IgG levels were similar amongst all adjuvants, while isotyping revealed slightly lower IgG1 levels in CAF10b-immunized mice and lower IgG2c levels in CAF01- and CAF09^hi^-immunized mice. All H107/CAF® vaccines induced vaccine-specific CD4 T cells, including IFNγ-producing (Th1) and IL-17A-producing (Th17) responses (Fig. 2B). CAF®01 and CAF®10b were the most effective at inducing vaccine-specific Th1/Th17 responses (Fig. 2C,D). In contrast, the CAF®09 formulations were less immunogenic, inducing significantly lower Th1/Th17 responses (Fig. 2C). To study the functional capacity of the immune responses, we next took advantage of the murine Mtb infection model, where a combined Th1/Th17 response is suggested optimal for protection (*23, 24, 32, 33*). Indeed, following aerosol challenge with Mtb, H107 adjuvanted with either CAF®01 or CAF®10b provided the greatest CFU reduction (2.53 log and 2.32 log, respectively, compared to control animals) (Fig. 2E).

**Figure 2.**
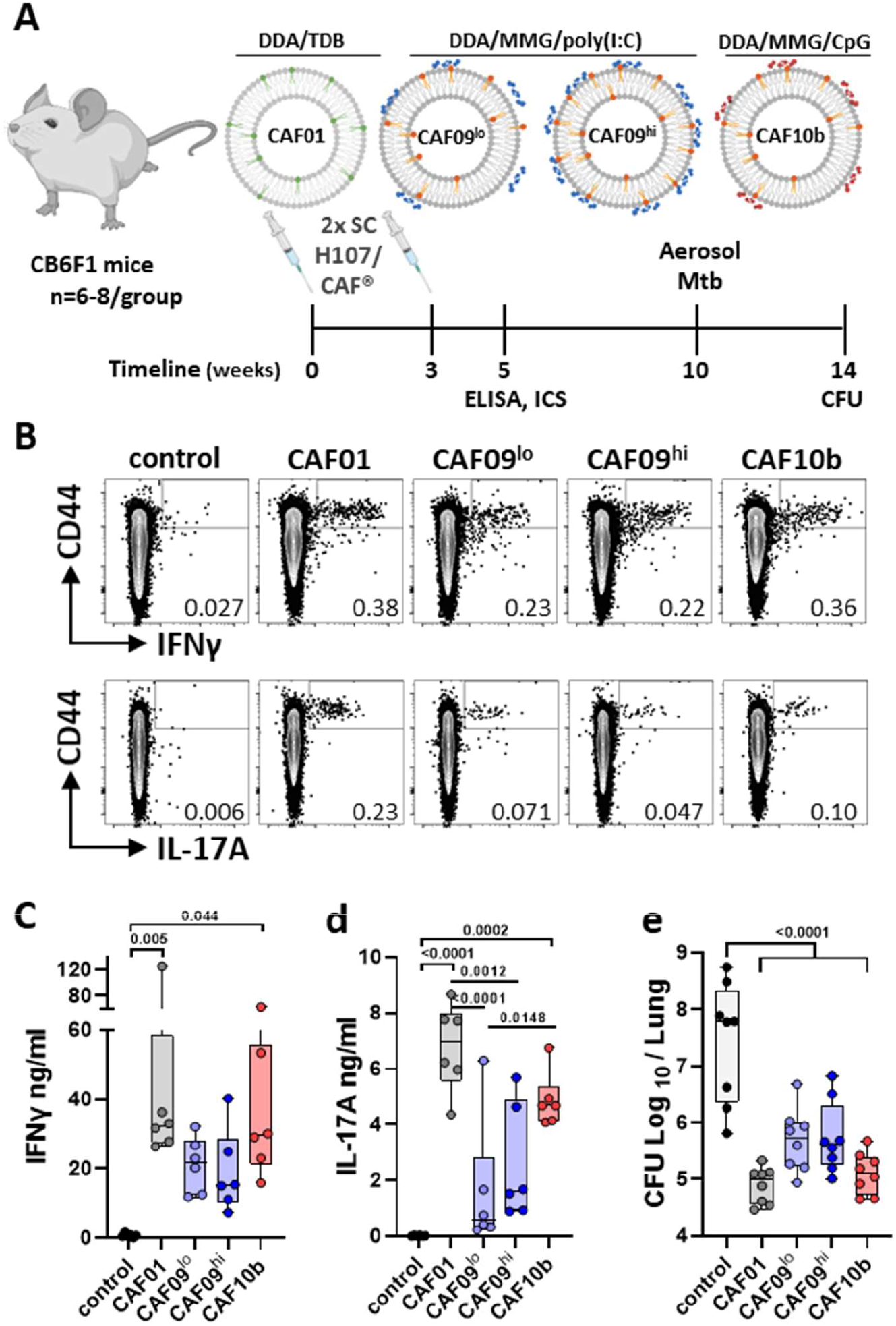
CAF® induction of Th1 and Th17 responses in mice. **(A)** CB6F1 mice were immunized with the H107 antigen adsorbed to different CAF® adjuvants as depicted. **(B-D)** Splenocytes taken 2 weeks after the last immunization were stimulated *ex vivo* with H107. (B) ICS analysis, showing representative dot plots of CD4 T cells following ICS assay from each vaccine group. Number shown depicts % cytokine positive CD4 T cells (live, lymphocytes, CD3+, CD8-, CD4+). (C,D) ELISAs for IFNγ (B) and IL-17A (C) secretion from H107-stimulated splenocytes. **(E)** Bacterial load in the lungs of control and H107/CAF®-immunized mice 4 weeks after aerosol Mtb infection. (C-E) Symbols display individual mice; Bars represent median, IQR, min. and max values. P values, one-way ANOVA with Tukey’s posttest.

In summary, CAF®10b induced similar Th1/Th17 responses as CAF®01 in mice and provided robust protection against murine Mtb aerosol challenge, when administered with the H107 antigen.

### CAF®10b induces robust antibody responses in non-human primates

To assess the translational potential of CAF®10b, we sought to directly compare specific cellular and humoral immunogenicity of previous CAF® platform adjuvants in NHPs. In parallel to the mouse studies, the H107 antigen was combined with either CAF®10b, CAF®01, or the ‘low’ or ‘high’ formulations of CAF®09 (Fig.3). CAF®01 served as a benchmark in these studies as it has previously been combined with several antigens, including the tuberculosis vaccine antigen H56 and the chlamydia antigen CTH522 in NHP immunization studies (*25, 34, 35*). The vaccines were given as a two-dose intramuscular immunization regimen with four weeks spacing in cynomolgus macaques (Fig. 3). CMI and humoral immune responses were determined via blood samples taken at baseline prior to immunization and at multiple time points after immunization, including 6 months after the final immunization (Fig. 3). Overall, all four CAF® adjuvants were well-tolerated; a transient increased body temperature on the day after each immunization, typically 0-2°C, was observed for immunized NHPs, but otherwise did not vary relative to controls (Fig. S4A). Similarly, overall weight changes were not different in the immunized groups (Fig. S4B), and complete blood counts taken bi-weekly over the study’s course did not show measurable differences in their white blood cellular composition (Fig. S4C,D).

**Figure 3.**
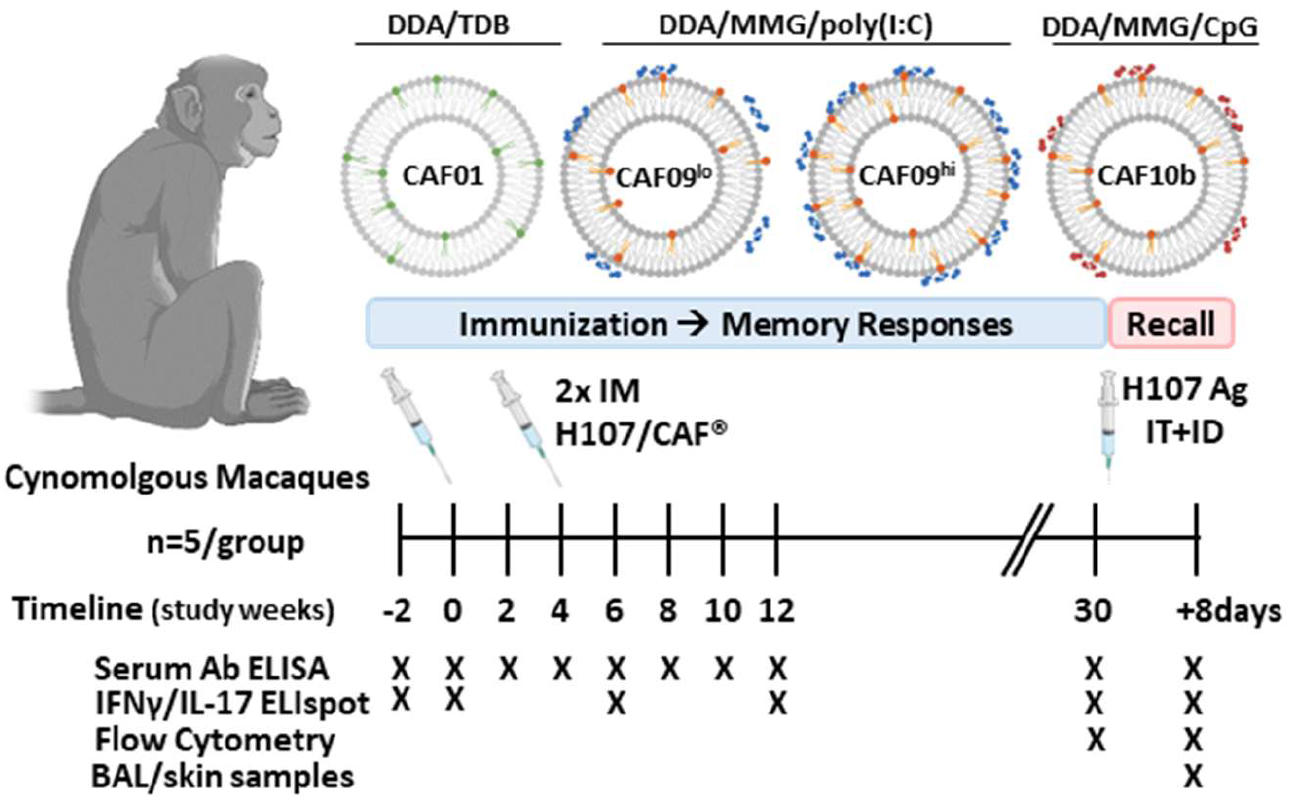
CAF® adjuvant comparison in Non-Human Primates. Cynomolgus macaques were immunized with the H107 antigen adsorbed to different CAF® adjuvants as depicted. Animals were monitored for baseline, initial, and memory immune responses in the blood (serum, PBMC). In the antigen recall phase, animals were administered non-adjuvanted H107 protein antigen into the lung and skin and the systemic (serum, PBMC) and local (skin, BAL) recall responses assessed.

All adjuvants induced H107-specific IgG in the serum of NHPs after the first immunization, which was further boosted after the second immunization (Fig. 4A). Peak anti-H107 IgG responses were achieved 2 weeks after the booster for all groups (wk6), wherein median H107/CAF®10b and H107/CAF09^hi^ induced greater than 2-fold higher IgG relative to H107/CAF®01 with increases of 0.323 log and 0.374 log arbitrary units, respectively (Fig. 4B). The median serum anti-H107 IgG levels induced by CAF®10b and CAF09^hi^ remained consistently higher than CAF®01 and CAF®09^lo^ at all time points investigated (Fig. 4A). Cumulative IgG responses confirmed induction of significant responses versus non-immunized control animals over the 30-week period of the study (Fig. 4C). Serum H107-specific IgA was detected after the second immunization in animals immunized with CAF®01, CAF®09^hi^, and CAF®10b (Fig. 4E,F). However, the kinetics of sustained antibody responses varied, and when integrated over 30 weeks, only H107 adjuvanted with CAF®09^hi^ and CAF®10b demonstrated a significant IgA response (Fig. 4F).

**Figure 4.**
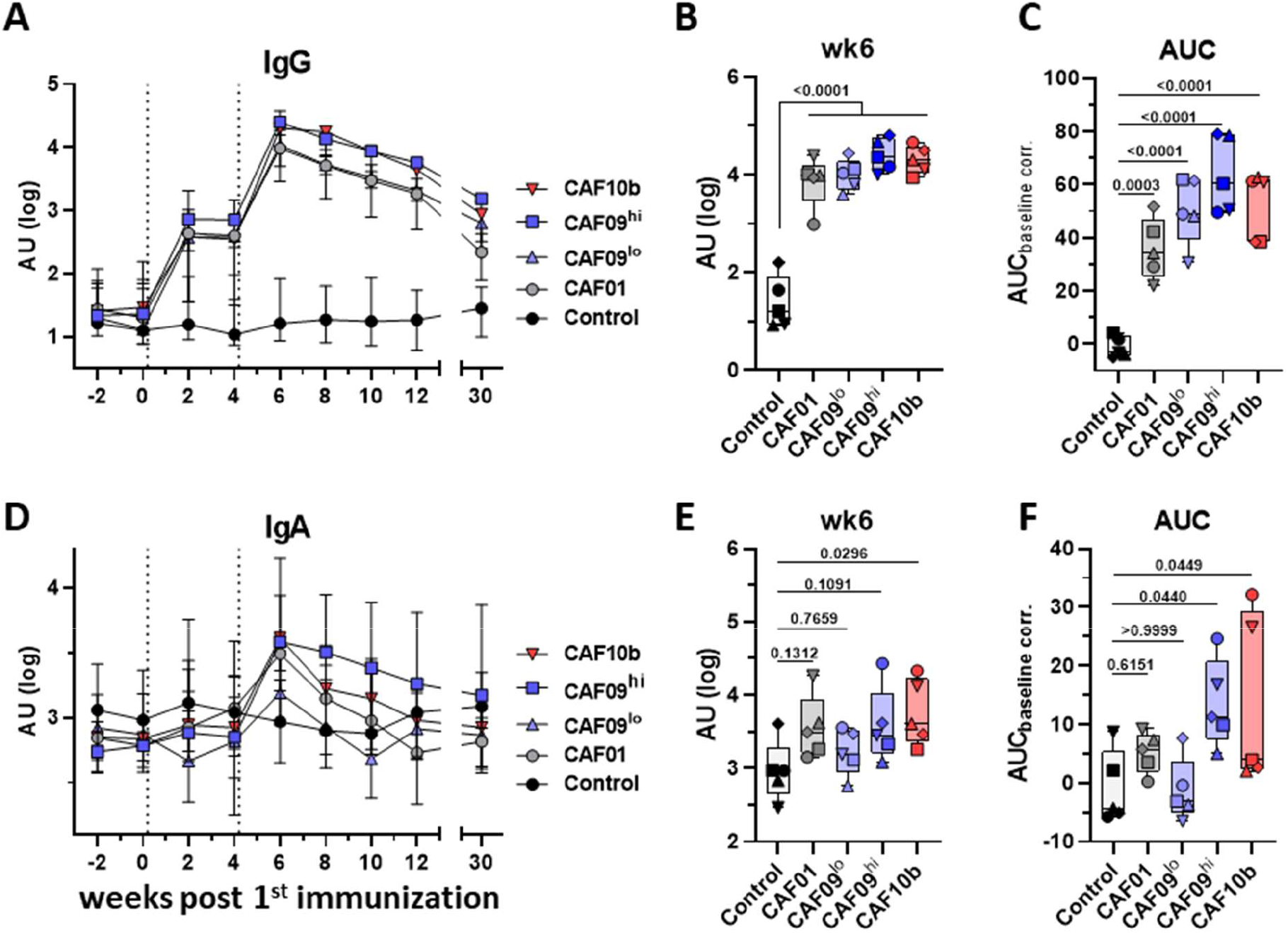
IgG and IgA responses are influenced by vaccine composition. H107-specific IgG **(A-C)** and IgA **(D-F)** antibody levels in the serum measured by ELISA before and after H107/CAF® immunization of Nonhuman primates. (A,D) Immunizations are indicated by dotted lines. Symbols represent median ±IQR. AU, arbitrary units. (B,E) Peak responses at week 6. (C,F) Responses integrated over the 30-week time course calculated as area under the curve after baseline subtraction (AUC) are depicted. (B,C,E,F) Bars represent median ± IQR, min. and max. Symbols, individual animals. P values from One-way ANOVA with Dunnett’s post-hoc test of log-transformed data.

Overall, all CAF® platform adjuvants investigated induced vaccine-specific antibody responses, with CAF®09^hi^ and CAF®10b displaying both the highest peak and sustained antigen-specific IgG and IgA responses.

### CAF®10b induces long-term Th1 and Th17 responses in NHPs

A major focus of this study was to investigate the CMI-inducing capacity of these adjuvants, including Th1 and Th17 responses. We prioritized sensitive IFNγ/IL-17 ELISpots over flow cytometry for the post-vaccination responses, for maximizing the chances of detecting Th17 responses that are rarely present in blood (*36, 37*). Therefore, we used stimulation of PBMCs in sensitive ELISpot assays to assess the induction of H107 vaccine-specific T cells producing IFNγ (Th1) and IL-17A (Th17) cells in the blood before and after immunization. Antigen stimulation revealed systemic IFNγ responses in all vaccine adjuvant groups, with the peak response detected two weeks after the final immunization (six weeks post first immunization), after which there was a contraction of responses (Fig. 5A). At peak, CAF®09^hi^ and CAF®10b showed the highest vaccine-specific IFNγ responses, with all animals (5/5) displaying spot-forming cells (SFC) counts above those of any control animal (Fig. 5A, right). Interestingly, at 30 weeks post first immunization, only the group of H107/CAF®10b-immunized animals had a significant detectable IFNγ ELISpot response, demonstrating sustained long-term CMI memory. Similar to IFNγ, IL-17A ELISpot results demonstrated vaccine-specific responses with similar kinetics, with individual animals in all immunization groups demonstrating vaccine specific IL-17A responses two weeks after the final immunization (Fig. 5B, left). However, 30 weeks post immunization, only the H107/CAF®10b group had a durable long-term Th17 memory response, with 4/5 animals still displaying IL-17A SFC counts above those of any control animal (Fig. 5B, right).

**Figure 5.**
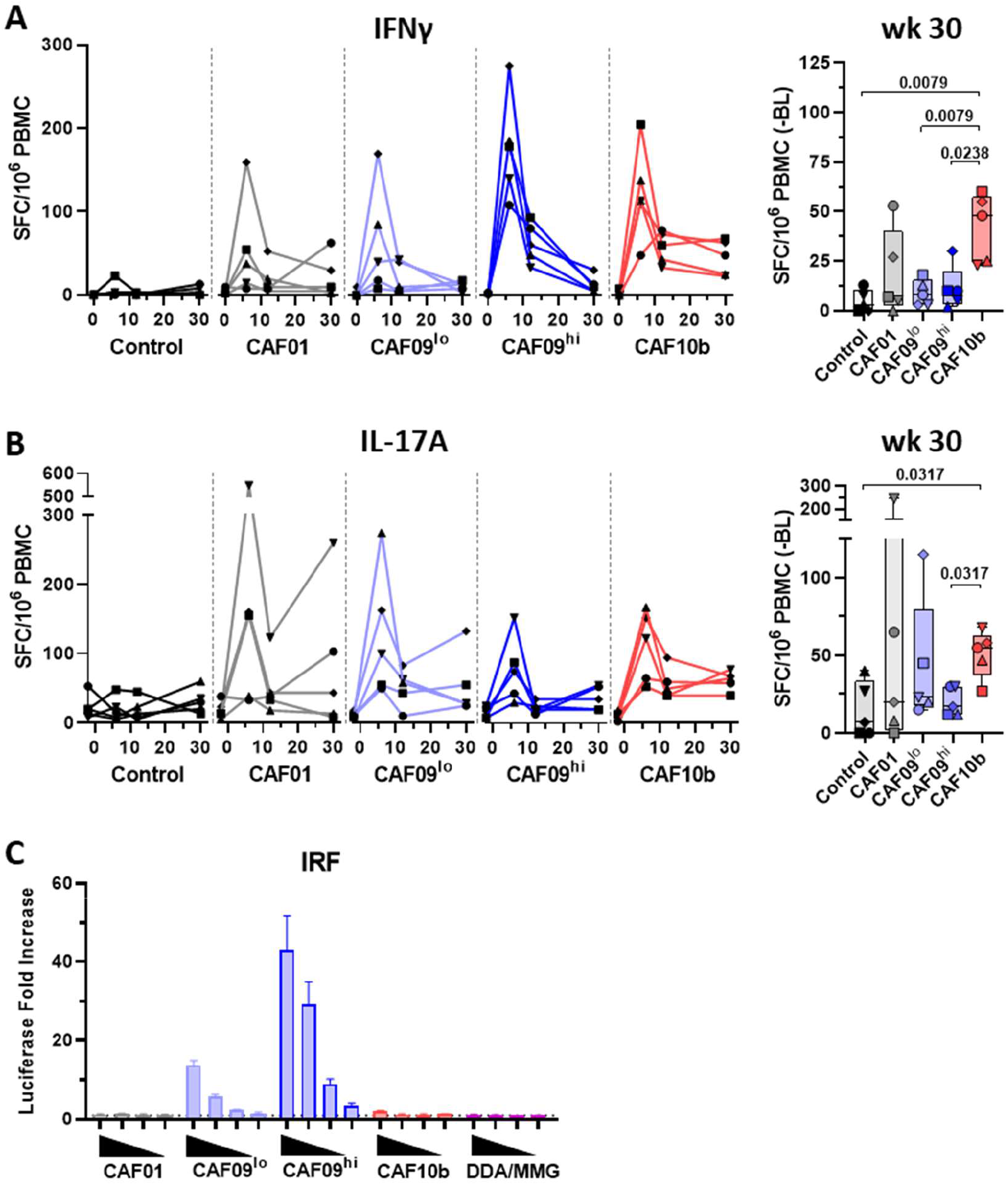
CAF® adjuvant induction of Th1 and Th17 responses differ in NHP. **(A,B)** PBMCs from immunized Non-human primates assessed by ELISpot before and after immunizations. PBMCs were stimulated *ex vivo* with for H107-specific (A) IFNγ and (B) IL-17A production. Symbols and lines indicate individual animals (left panels). Memory responses at 30 weeks post first immunization corrected for pre-immunization baseline SFC and plotted for each H107/CAF® immunized group (A,B right panels). Bars represent median ± IQR, min. and max. Symbols, individual animals. P values shown from Mann-Whitney U tests. Kruskal-Wallis test (A) p=0.038 (B) p=0.1253 **(C)** PMA+Ionomycin-differentiated THP1-Dual™ cells stimulated overnight with serial 2-fold dilutions for various CAF® liposome formulations, from 25 to 3.125 μg/mL DDA, were assessed for induction of IRF-driven secreted luciferase as a fold increase versus untreated cells. Bars, mean±SD of triplicates, representative data from two similar experiments.

We noted that CAF09^hi^ induced the highest initial vaccine-specific IFNγ response, accompanied by a low, transient IL-17A response. In contrast, CAF®10b was able to induce both sustained IFNγ and IL-17A responses. Type I interferons can have inhibitory effects on Th17 induction (*38–41*) and TLR3 signals via TIR-domain-containing adapter-inducing interferon-β (TRIF) (*42*). Therefore, we hypothesized that induction of the type I interferon pathways by Poly(I:C) could be responsible for the diminished Th17 response of the CAF®09^hi^ formulation. Indeed, stimulation of human monocytic THP-1 cells with an integrated type I interferon response factor (IRF)-driven luciferase reporter (THP1-Dual cells) with CAF®01, CAF®10b, or DDA/MMG liposomes alone did not induce type I interferons (Fig. 5C). In contrast, treatment with CAF®09 liposomes induced IRF-elements in a Poly(I:C) dose-dependent manner (*i.e*. CAF®09^hi^ > CAF®09^lo^, Fig. 5C), consistent with the suppression of Th17 responses observed *in vivo* (Fig. 5B).

In summary, intramuscular immunization of NHPs with CAF®10b led to robust induction of both Th1 and Th17 responses. These responses were detected in circulation at least 6 months after intramuscular administration.

### *In vivo* antigen delivery recalls local and systemic memory responses

Finally, to further assess the quality, capacity, and potential of the vaccine-induced adaptive memory responses, we investigated local and systemic recall of CMI and antibody responses *in vivo* after antigen recall. Six months after the final immunization, H107 protein and an irrelevant control protein (CTH522 (*43*)) were given intradermally at distal sites on each animal’s upper back. Concurrently, H107 was given into the airways via bronchoscope instillation into the caudal right lung. Skin sites were monitored for delayed-type hypersensitivity (DTH) responses, where only a few immunized animals displayed minimal local redness and light swelling at the H107 injection sites (Fig. S5A). In contrast, sequential ^18^F-Fluorodesoxyglucose (^18^F-FDG) positron emission tomography with computed tomography (PET-CT) scanning of selected animals before and after instillation demonstrated pulmonary recall responses in the right lungs of H107/CAF®10b-immunized NHP, but not in contralateral lungs or in control animals (Fig. 6B). Such responses were detectable as early as three days after instillation and were resolved by day seven post recall, demonstrating a short-lived local inflammatory response to antigen recall (Fig. 6B, Fig. S5B). Despite the intradermal antigen injection sites also being within the PET-CT scanning area, no local skin inflammatory responses were detected in these animals, consistent with a lack of observed DTH (Fig. S5A).

**Figure 6.**
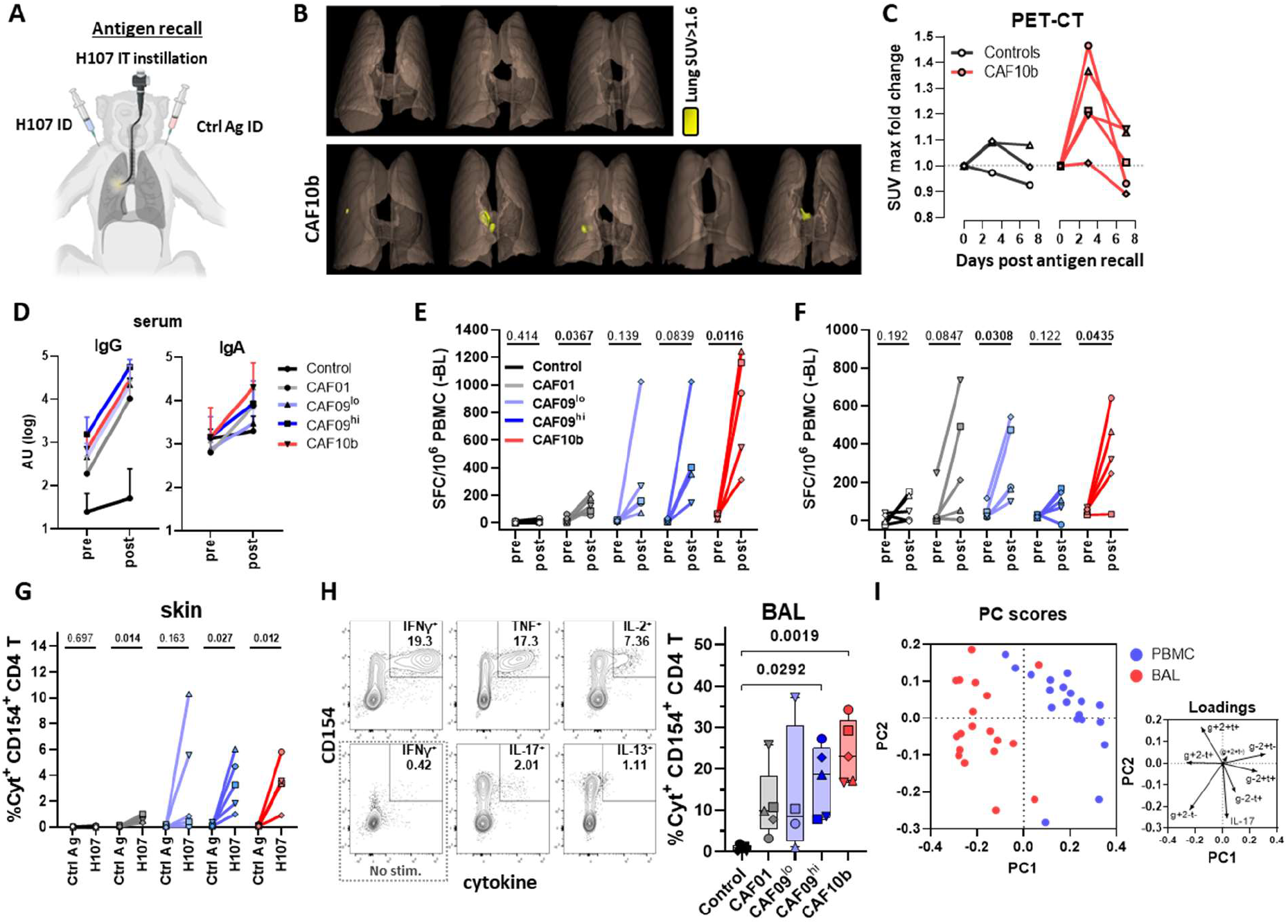
*In vivo* antigen delivery recalls local and systemic memory responses. Six months after the final immunization, non-human primates were given an *in vivo* antigen recall in the skin and lungs with H107, or a control protein, as shown in **(A)**. **(B)** PET-CT scans of the selected Control (n=3) and H107/CAF®10b-immunized (n=5) animals depicting local lung inflammatory responses 3 days after H107 antigen recall, as standardized uptake value (SUV) > 1.6. **(C)** The SUV maximum fold change relative to pre-recall (day 0) for the antigen-receiving lung lobes at 3 and 7 days after antigen recall for the NHPs assessed by PET-CT. **(D)** Serum H107-specific IgG (left) and IgA (right) antibody levels as measured by ELISA before (pre) and post *in vivo* antigen recall of H107/CAF®-immunized Non-human primates. Symbols represent mean±SD. AU, arbitrary units. **(E,F)** PBMCs from control and H107/CAF®-immunized NHPs were assessed by ELISpot assay pre- and post *in vivo* antigen recall. PBMCs were stimulated *ex vivo* for H107-specific (E) IFNγ and (F) IL-17A production. **(G)** The percentage of CD4 T cells isolated from skin biopsies collected from control antigen (Ctrl Ag) and H107 ID injection sites that express IFNγ, TNFα, IL-2, IL-17, and/or IL-13 after *ex vivo* stimulation with H107 protein, as assessed by intracellular cytokine staining (ICS). (E-G) Symbols and lines, individual animals. P values within groups from paired T test. **(H)** Sample contour plots of ICS analysis of CD4 T cells isolated from the BAL of a representative H107/CAF®10b-immunized animal 8 days after *in vivo* antigen (left), and the resultant summary data of the total % H107-specific cytokine-expressing BAL CD4 T cells. Bars represent median ± IQR, min. and max. Symbols, individual animals. P values from Mann-Whitney U tests. Kruskal-Wallis test p=0.0097 **(I)** Principle component analysis (PCA) of H107-specific CD4 T cells from BAL (red) and PBMC (blue) for IL-17 and combinatorial IFNγ/TNFα/IL-2 expression. PC1 and PC2 loading axes shown. Symbol, individual H107/CAF®-immunized animals.

Eight days after antigen administration, all vaccine groups were assessed for local (skin/lung) and systemic recall responses. Antigen recall increased serum H107-specific IgG significantly in all vaccine groups, demonstrating a systemic booster response to antigen recall (Fig. 6C). Serum H107-specific IgA was also increased in the groups having received CAF®01, CAF®10b and CAF®09^hi^, but not CAF®09^lo^ (Fig. 6C), consistent with the pattern of antigen-specific IgA after immunization (Fig. 4b). In contrast, vaccine-specific IgG, but not IgA, was detected in bronchoalveolar lavage (BAL) fluid of all H107/CAF®-immunized groups compared to controls (Fig. S5C). Vaccine-specific antibodies were also assayed locally in the skin, however equal H107-specfic IgG levels were detected in the biopsies taken at both the H107 and control protein sites, and therefore likely only reflected the systemic serum levels.

Systemic vaccine-specific CMI responses were also highly expanded by *in vivo* antigen recall. Notably, ELISpot analysis of PBMCs demonstrated a significant 19.6-fold median expansion (interquartile range 11-50, p=0.0116) in H107-specific IFNγ-producing (Th1) cells in H107/CAF®10b-immunized NHPs (Fig. 6E). IL-17A-secreting (Th17) cells were also greatly expanded in H107/CAF®10b animals (p=0.0435) confirming the presence of recallable memory Th17 cells following immunization (Fig. 6F). Systemic memory Th1 and Th17 responses were also expanded, in H107/CAF®01 and H107/CAF®09^lo^ animals. However, consistent with the lack of Th17 cells after initial H107/CAF®09^hi^ immunization (Fig. 5B), only Th1 responses were expanded in H107/CAF®09^hi^ animals (Fig. 6F).

Comparing the skin-infiltrating cells at the H107 and control protein sites of injection by flow cytometric analysis demonstrated vaccine-specific CD4 T cell recall responses (Fig. 6G). *Ex vivo* H107 stimulation of skin biopsy-derived cells followed by intracellular cytokine staining (ICS) for production of IFNγ, TNFα, IL-2, IL-17, and/or IL-13 (Fig. 6G) showed a specific recruitment of H107-specific CD4 T cells to only the H107 injection site for all vaccine groups, with the highest frequency of H107-specific CD4 T cells found in the H107/CAF®09hi and H107/CAF®10b cohorts (Fig. 6G, for gating strategy see Fig. S6A). Local pulmonary cellular recall was also assessed by ICS of cells collected from BAL of the right lungs eight days after antigen instillation. Here as well, CD4 T cells specific for H107 were found in all immunized groups, with the percentage of H107-specific CD4 T cells highest for those previously immunized with H107/CAF®10b (Fig. 6H). Upon H107 restimulation *ex vivo*, the primary CD4 T cell-produced cytokines observed in the BAL were the Th1 cytokine IFNγ, TNFα and IL-2, with a notable population of IL-17-producing Th17 cells and a lesser population producing IL-13 (Fig. 6H). Notably, H107-specific cytokine production by ICS was not observed for CD8 T cells in the skin, BAL, or PBMC, supporting that the antigen-specific CMI responses induced by these CAF®-adjuvated H107 vaccines and measured by ELISpot and ICS were Th1/Th17 CD4 T cells (Fig. 6G,H; Fig. S6B-D). We found no significant correlation between H107-specific responses in the blood prior to antigen recall and subsequent CD4 T cell responses in the BAL, suggesting that the systemic CD4 T cell memory does not predict *in vivo* recall in the lung (Fig. S6E). Instead, there was a positive correlation between H107-specific CD4 T cells in the blood and BAL *after* recall, indicating that local and systemic recall responses are related (r=0.5748, p=0.013, Fig. 6E). Finally, a combined Boolean gating and principle component analysis (PCA) of cytokine expression profiles of these post-recall H107-specific CD4 T cells, demonstrated a site-specific phenotypic distinction between PBMCs and BAL (Fig. 6I). The biplot loadings showed that while the PBMC were differentiated by TNF+IL-2+ and IL-2+ CD4 T cells, the BAL cells were characterized by IFNγ+ and IFNγ+TNF+ CD4 T cells, reflecting an effector/effector memory phenotype (*44*).

In summary, the *in vivo* antigen administration 6 months after immunization demonstrated robust systemic and local recall responses at the sites of instillation revealing the relative memory CD4 T cell response of each CAF® vaccine formulation. Overall, CAF®10b displayed the highest recall responses with significant expansion of both Th1 and Th17 cells.

## DISCUSSION

Difficult-to-target diseases, for which conventional approaches are incomplete or have failed, pose special challenges to vaccine design. Thus, development of improved vaccines against such targets, likely depends on vaccine platforms with diverse immune signatures. Since the delineation of Th1 and Th2 cells in the 1980s (*45*), several other T cell subsets have been identified, including Th17 cells that were described in 2005 (*46, 47*). Now, almost two decades later, current licensed adjuvants induce antibodies, Th1 and Th2 cells, but not Th17 cells, despite their implication in protection against major pathogens like influenza (*10, 12*), chlamydia (*48*), Klebsiella pneumoniae (*49*), group A streptococci (*50, 51*) and tuberculosis (*5, 52*). Notably, in experimental settings, immunizations that induce Th17 responses have been limited to mucosal delivery, including for live viral vectored vaccines and BCG (Bacillus Calmette Guérin) (*7, 53–55*). With the goal of closing this gap, our study introduces CAF®10b as a new liposomal adjuvant that induces strong humoral and cellular responses in both mice and NHPs with robust Th1/Th17 memory after intramuscular injection. Consisting of all-synthetic components, this adjuvant is compatible with large-scale manufacturing from readily available substrates, mitigating potential availability issues as seen with some of the existing adjuvants derived from natural sources (*13, 14*).

Inspired by data from the mouse model (*27*), we initiated our search for an improved adjuvant for human use by assessing the potential of CpG2006 as an add-on to the CAF® platform (*15*). Consistent with previous results (*27, 56*), we confirmed that CpG2006 activates human TLR9 and that highly stable CAF®10b liposomes could be formed by formulating CpG2006 with DDA/MMG. Although CpG alone did not induce Th17 cells, we found that CpG2006 activated murine TLR9 and that the incorporation of CpG2006 within CAF®10b liposomes increased the CAF®-driven Th1/Th17 responses in mice, while simultaneously greatly reducing the systemic inflammatory effects of CpG2006 alone. When combined with the TB vaccine antigen H107 (*23*), this CAF®10b-induced Th1/Th17 phenotype was associated with increased protection against murine Mtb challenge compared to CAF09® formulations (*26, 31*) that induced a more strict Th1 response. While this demonstrates the functional capacity of CAF®10b-induced immunity to protect against an airway pathogen, it also adds to the accumulating support for a protective role of vaccine-induced Th17 cells against TB (*5, 7, 23, 52, 53, 55*).

To better assess the translational potential of CAF®10b for human use, we turned to the NHP model and compared the specific humoral and cellular responses to previous CAF® formulations, CAF®01 and CAF®09. Here we observed that two intramuscular injections with H107/CAF®10b led to strong serum IgG responses that were sustained at approximately 2-fold (0.323 log) higher than for CAF®01. This is remarkable, as CAF®01 has already been shown to induce 5-6 fold higher IgG titers than alum in a head-to-head clinical trial (*17*). In addition, we also observed induction of serum IgA responses, which could be important for certain viral targets, as serum IgA has shown superior capacity to neutralize SARS-CoV-2 virus (*57, 58*).

While antibodies play an important role in immunity against many pathogens, control of some disease targets also benefits from a potent cellular Th1 response. These include bacterial and viral pathogens like chlamydia (*59*), TB (*24*), SARS-CoV-2 (*60*), RSV (*61, 62*), and influenza (*63*) where effective pathogen control is associated with a Th1 phenotype, in contrast to Th2 cells that associate with disease severity (*60, 63–66*). We were therefore intrigued to find that CAF®10b induced a robust Th1 memory response that was detectable in circulation six months after intramuscular injection. Interestingly, CAF®09^hi^ induced slightly higher peak Th1 responses, but after six months, this had contracted to baseline levels, highlighting that peak responses do not necessarily predict systemic memory. This is conflicting with models where the cellular memory pool is formed as a subset of the effector response, which has been described for effector memory CD4 T cells (*67*) and memory CD8 T cells (*68, 69*). However, central memory CD4 T cells are generated as an independent subpopulation of the effector response (*67*) and the discrepancy between peak and memory responses could therefore be explained by differential imprinting of the central memory T cell pool. Although our study was not designed to answer this question, our findings stress the importance of performing long-term experiments to assess true vaccine memory.

In addition to Th1 cells, we also found a substantial and persistent Th17 response in NHPs vaccinated with CAF®10b, and to a lesser degree with CAF®01 and CAF®09b This is to our knowledge the first report of an adjuvant that induces a robust Th17 response in NHPs after intramuscular administration. Intriguingly, the IL-17 response for CAF®10b was detectable at termination of the study (26 weeks after the last immunization), which is consistent with previous data from the mouse model, showing that vaccine-induced Th17 cells are maintained as a phenotypically stable subset (*70*). In contrast to that observed with CAF®10b, we did not find detectable Th17 responses for CAF®09^hi^. CAF® liposomes, containing either TDB or MMG, are described to drive Th1/Th17 responses via stimulation of the C type lectin receptor, Mincle, which leads to induction of proinflammatory cytokines like IL-1β, IL-6, TNF-α and TGF-β (*15*). However, a recent report showed that CAF®09 administration drives expression of type-I IFN related genes in mice (*71*), and in this study we also observed that CAF®09 activated IRF in human THP-1 reporter cells in a dose dependent manner (CAF®09^hi^ > CAF®09^), whereas CAF®01 and CAF®10b had no effect. These data suggest that poly(I:C), included in CAF®09, suppresses Th17 responses through induction of type I IFNs, in a dose dependent manner, as has been described in settings of autoimmunity and bacterial infection (*39–41*). Recognizing that Th17 cells have stem cell-like features, with some studies reporting superior long-term survival capacity over Th1 cells (*72–74*), this could also be part of the explanation for why CAF®10b induced greater T cell memory responses than CAF®09^hi^, despite showing similar peak Th1 responses.

Effective vaccine-memory is characterized by the capacity to mount a rapid recall response to challenge in relevant tissues. As a functional approach to studying immune memory, we applied an *in vivo* antigen recall model to investigate *antigenic challenge*, where unadjuvanted antigen was delivered into the skin and lungs of NHPs six months after vaccination. Here we observed significant recall responses systemically and locally, including dramatically increased H107-specific IgG in the serum and presence in the lungs (BAL) for all adjuvants tested. Blood CMI responses were also dramatically expanded for all vaccine groups. Local CMI responses were specifically localized to H107-injected skin sites, and H107-specific CD4 T cells were found in the BAL of all immunized groups (although with lesser statistical confidence for CAF®01 and CAF®09^lo^ versus control). Interestingly, *in vivo* antigen recall responses were evident for CAF®01 and CAF®09-immunized animals even though such CMI memory responses were undetectable in PBMCs 30 weeks after immunization, demonstrating the sensitivity of the recall assay for identifying subtle memory responses. Notably, the relative hierarchy of the given responses (*e.g*. IgG, Th1, Th17) was not changed by the antigen recall, i.e. CAF®10b had the highest memory Th17 response pre- and post-recall, while CAF®09^hi^-immunized animals did not have a significant Th17 response above control animals. In general, we found the strongest systemic and local CMI recall responses in the CAF®10b-adjuvanted vaccine group, although the antibody responses were similar to those observed with the CAF®09^hi^-adjuvanted vaccine.

Of specific interest in this study, we observed a striking expansion of Th17 cells after antigen recall, measured by both ELISpot and flow cytometry. This is in line with mouse data for CAF®01 showing that systemic vaccination leads to a small pool of Th17 cells that significantly expand in response to mucosal infection by either vaginal *Chlamydia trachomatis* (*48*) or aerosol Mtb (*75*). While these data form conclusive evidence that vaccine-induced Th17 cells respond to local antigen recognition in NHPs, they also highlight the antigen recall approach as a sensitive means to amplify and detect Th17 cells that can otherwise can be underrepresented in blood samples (*36, 37*). We were therefore intrigued to see that the inflammation induced by the antigen recall was highly localized and short-lived (<7 days), which supports the use of this approach in humans to study local vaccine responses. Such studies could focus on lung sampling, similar to experimental studies investigating pulmonary BCG instillation (*76*), or sampling of the upper airway mucosa, as has recently been used to study local responses to SARS-CoV-2 (*77*). The latter approach being a non-invasive alternative to performing multiple bronchoscopies, would likely be more feasible in early clinical studies of new vaccine candidates or larger head-to-head studies comparing different vaccine candidates.

Head-to-head studies are costly to perform, even in the pre-clinical setting. This also explains why most adjuvant studies are performed in mice. However, in this study we conducted parallel head-to-head experiments in both mice and NHPs, observing notably different hierarchies of the adjuvant-mediated responses. For example, while the Th1/Th17 responses induced by CAF®01 were relatively the greater than the CAF®09^lo/hi^ in mice, this was not so in NHPs, indicating differences in the immune recognition between these two species. Similarly, CAF®09 has previously been shown to induce greater cellular and humoral responses than CAF®01 in cattle (*78*). These species-specific differences could in part be driven by differential expression of pattern recognition receptors and/or differential sensitivity to different immunostimulants, as has been reported for Mincle-ligands (*79*) and CpGs (*80*). Therefore, even though we observed that some aspects of the adjuvant responses were consistent across mice and NHPs (*e.g*. the diminished Th17 responses of CAF®09), the species-specificity of our data underscores the importance of broad species testing of vaccines in preclinical settings and when evaluating candidates for further clinical development. In that regard, the relative consistency of CAF®10b to induce robust Th1 and Th17 responses in mice and NHP maybe prove a benefit in future translations studies across these species.

Overall, we have developed a new liposomal adjuvant, CAF®10b, that is optimized for human immunogenicity by stabile incorporation of an agonist that specifically activates human TLR9. With strong antibody responses, robust Th1/Th17 memory induction after intramuscular injection, and significant mucosal recall responses in NHPs, CAF®10b has inherent potential to fill an important gap in the existing pipeline of vaccine adjuvants. Based on these properties, we have initiated GMP manufacturing of CAF®10b in preparation for clinical testing.

## MATERIALS AND METHODS

### Antigen and Adjuvant production and characterization

The H107 fusion protein is composed of eight Mtb antigens, of which ESAT-6 is repeated four times: PPE68-[ESAT-6]-EspI-[ESAT-6]-EspC- [ESAT-6]-EspA-[ESAT-6]-MPT64-MPT70-MPT83. Recombinant H107 protein was produced and purified as in accordance with established methods (*23*). Briefly, the DNA construct was codon-optimized for expression in *E. coli* and inserted into the pJ411 expression vector (ATUM, Menlo Park, CA, US). H107 contained a His-tag at the N-terminal end (MHHHHHH-). After transformation into *E. coli* BL21 (DE3) (Agilent, DK), protein expression was induced with 1 mM isopropyl ß-d-1-thiogalactopyranoside in 3-liter cultures, and the H107 protein was purified from inclusion bodies by metal chelate chromatography followed by anion-exchange chromatography. Preparations were sterile filtered in 20mM Glycine, pH9.2.

Control recall protein, CTH522 (*43*) (composed of *Chlamydia trachomatis-specific* antigens) was produced and supplied by Vaccine Development at Statens Serum Institut according to GMP in 20 mM Glycine, pH 9.2 as previously described (*17*).

CAF® adjuvants were prepared as previously described for CAF®01 and CAF®09^lo^ (CAF®09b) (*81*). CAF09^hi^ was produced analogously, but with double concentration of MMG and Poly(I:C). CAF®10b was produced as previously described (*27*), but using CpG ODN 2006 (InVivoGen or BianoScience, Gera, Germany) with the sequence 5’-tcgtcgttttgtcgttttgtcgtt-3.

Adjuvants were characterized for particle size and polydispersity index (PDI) by dynamic light scattering (photon correlation spectroscopy technique). Surface charge was analyzed by measuring the zeta potential (laser-Doppler electrophoresis). For the size measurements, the samples were diluted 10 times, whereas for the zeta potential measurements, the samples were diluted 100 times in milli-Q water. The measurements were performed at 25°C by using a Zetasizer Nano ZS (Malvern Instruments, Worcestershire, UK) equipped with a 633 nm laser and 173° detection optics. Malvern DTS v.6.20 software was used for data acquisition and analysis. DDA and MMG content was measured by HPLC using an Ultimate 3000 HPLC with a Corona Veo Charged Aerosol Detector (Thermo Fisher Scientific, Waltham, MA, USA). Chromeleon software version 7.2 SR5 was used for data acquisition and analysis. CpG content was measured in a 96 well plate using Ribogreen™ fluorescence staining kit from Thermo Fisher and a SpectraMax i3 plate reader (Molecular devices, San Jose, CA, USA). CpG content was measured in a 96 well plate using Ribogreen™ fluorescence staining kit from Thermo Fisher and a SpectraMax i3 plate reader (Molecular devices, San Jose, CA, USA). Differential Scanning Calorimetry Analysis of the liposomes was determined using a MicroCal PEAQ-DSC version 1.61 (Malvern Panalytical Nordic AB, Naerum, Denmark) for data acquisition and analysis. Stability was measured for up to 9 months.

### Mouse studies

#### Mice

Six-to-eight week old female CB6F1 (H2^b,d^) mice were obtained from Envigo (Netherlands). Mice were randomly assigned to cages upon arrival and acclimatized at least one week to the animal facility. During the course of the experiment, mice had access to irradiated Teklad Global 16% Protein Rodent Diet (Envigo, 2916C) and water ad libitum. Mice were housed under Biosafety Level (BSL) II or III conditions in individually ventilated cages (Scanbur, Denmark) and had access to nesting material (enviro-dri and soft paper wool; Brogaarden) as well as enrichment (aspen bricks, paper house, corn, seeds, and nuts; Brogaarden). Statens Serum Institut’s Animal Care and Use Committee approved all experimental procedures and protocols. All experiments were conducted in accordance with the regulations put forward by the Danish Ministry of Justice and Animal Protection Committees under license permit no. 2019-15-0201-00309 and in compliance with the European Union Directive 2010/63 EU.

#### Immunizations

Mice were immunized subcutaneously (SC) at the base of the tail two times with three-week intervals in a volume of 200 μl. Recombinant antigen was diluted in Tris-HCL buffer + 2% glycerol (pH 7.2) to a concentration of 2 μg and formulated in CCAF®01 (250/50μg DDA/TDB), CAF®10b(250/50/20μg DDA/MMG/CpG2006), CAF®09^lo^ (250/50/12.5μg DDA/MMG/polyI:C), or CAF®09^hi^ (250/100/25μg DDA/MMG/polyI:C).

#### Preparation of single-cell suspensions

Spleens were aseptically harvested from euthanized mice. forced through 70-μm cell strainers (BD) with the plunger from a 3 mL syringe (BD). Cells were washed twice in cold RPMI or PBS followed by 5 minutes centrifugation at 1800 rpm. Cells were finally resuspended in supplemented RPMI medium containing 10% fetal calf serum (FCS) as previously described (*34*). Cells were counted using an automatic Nucleocounter (Chemotec) and cell suspensions were adjusted to 2×10^5^ cells/well for ELISA and 1-2×10^6^ cells/well for flow cytometry. and serum collection.

#### Cytokine ELISAs

Splenocytes were cultured in the presence of 2 μg/mL recombinant protein antigen for 3 days at 37°C. Supernatants were harvested and analyzed for levels of IFNγ and IL-17A using enzyme-linked immuno-sorbent assay (ELISA) as previously described (*70*). Briefly, Microtiter plates (96-well; Maxisorb; Nunc) were coated with 1 μg/mL capture antibodies (IFNγ: clone R4-6A2, BD Pharmingen; or IL-17A: clone TC11-18H10.1, Biolegend) diluted in carbonate buffer. Free binding sites were blocked with 2% (w/v) skimmed milk powder (Natur Drogeriet, Matas, Denmark) in PBS. Culture supernatants were diluted in PBS with 2% Bovine Serum Albumin (BSA, Sigma-Aldrich) incubated overnight in plates. IFNγ was subsequently detected using a 0.1 μg/mL biotinylated rat anti-murine Ab (clone XMG1.2; BD Pharmingen) and recombinant IFNγ (BD Pharmingen) as a standard. IL-17A was detected using 0.25 μg/mL biotinylated anti-mouse IL-17A (BioLegend, clone: TC11-8H4) and recombinant IL-17A (BioLegend). Streptavidin HRP (BD Pharmingen, CA, US) diluted 1:5000 in PBS 1% BSA was used to detect bound biotinylated detection antibodies. The enzyme reaction was developed with 3,3’,5,5’-tetramethylbenzidine, hydrogen peroxide (TMB Plus; Kementec), stopped with 0.2 M H_2_SO_4_ solution and plates read at 450 nm with 620 nm background correction using an ELISA reader (Tecan Sunrise).

#### Intracellular Cytokine Staining (ICS)

Splenocytes stimulated *ex vivo* with 2 μg/mL H107 protein antigen in the presence of 1 μg/mL anti-CD28 (clone 37.51) and anti-CD49d (clone 9C10-MFR4.B) for 1 hour at 37°C, 5% CO_2_ followed by the addition of Brefeldin A to 10 μg/mL (Sigma Aldrich; B7651-5mg) and 5 hours of additional incubation at 37°C, after which the cells were kept at 4°C until staining. Cells were stained with surface markers diluted in 50% brilliant stain buffer (BD Horizon; 566349) using anti-CD4-BV510, anti-CD8-PerCP-Cy5.5, anti-CD44-BV786 at 4°C for 20min, before fixation and permeabilization using the Cytofix/Cytoperm kit (BD Biosciences) as per manufacturer’s instructions, followed by intracellular staining for anti-CD3-BV650, anti-IFNγ-PE-Cy7, anti-IL-2-APC-Cy7, anti-TNF-PE, anti-IL17-BV421. Cells were characterized using a BD LSRFortessa and the FSC files were manually gated with FlowJo v10 (TreeStar). Non-stimulated cells were used to set gate boundaries for cytokine markers.

#### Serum Collection and Assays

Blood was collected via cardiac puncture and serum isolated by centrifugation.

H107-specific antibody detection was performed by ELISA. Maxisorb Plates (Nunc) were coated overnight at 4°C with H107 (0.1 μg/ml) and then blocked for with 2% BSA in PBS for 2hrs at room temperature. Sera were diluted 30X followed by three-fold serial dilutions and incubated 2hr at room temperature 4°C on plates. Total IgG was detected with horse radish peroxidase (HRP)-conjugated goat anti-mouse IgG (Invitrogen) diluted 1:32000. IgG1 was detected with HRP-conjugated goat anti-mouse IgG1 (Southern Biotech) diluted 1:16000. IgG2c was detected with HRP-conjugated rabbit anti-mouse IgG2c (Southern Biotech) diluted 1:5000. The enzyme reaction was developed with 3,3’,5,5’-tetramethylbenzidine, hydrogen peroxide (TMB Plus; Kementec), stopped with 0.2 M H_2_SO_4_ solution and plates read at 450 nm with 620 nm background correction using an ELISA reader (Tecan Sunrise).

The Mouse U-plex kit for the cytokines IL-12p70, IL-6, TNFα, MCP-1 was performed according to the manufacturer’s instructions (Meso Scale Discovery) to measure cytokine concentrations in serum. The plates were read on the Sector Imager 2400 system (Meso Scale Discovery) and calculation of cytokine concentrations in unknown samples was determined by 4-parameter logistic non-linear regression analysis of the standard curve.

#### Aerosol infection and bacterial counting

Ten weeks after the first immunization, mice were challenged with Mtb Erdman (ATCC 35801 / TMC107). Mtb Erdman was cultured in Difco™ Middlebrook 7H9 (BD) supplemented with 10% BBL ™ Middlebrook ADC Enrichment (BD) for two-three weeks using an orbital shaker (~110 rpm, 37°C). Bacteria were harvested in log phase and stored at −80°C until use. Bacterial stocks were thawed, sonicated for five minutes, resuspended with a 27G needle, and mixed with PBS to the desired inoculum dose. Using a Biaera exposure system controlled via AeroMP software, mice were challenged by the aerosol route with virulent Mtb Erdman in a dose equivalent to 50-100 CFUs. To determine vaccine efficacy, Mtb CFU were enumerated in lungs of aerosol infected mice. Left lung lobes were homogenized in 3 mL MilliQ water containing PANTA™ Antibiotic Mixture (BD, cat.no. #245114) using GentleMACS M-tubes (Miltenyi Biotec). Lymph nodes were forced through 70-μm cell strainers (BD Biosciences) in 1 mL PANTA solution. Tissue homogenates were serially diluted, plated onto 7H11 plates (BD), and grown for 14 days at 37°C and 5% CO2. CFU data were log-transformed before analyses.

### Non-Human Primate (NHP) studies

#### NHPs

Cynomolgus macaques (*Macaca fascicularis*), aged 27-30 months (16 females and 14 males) and originating from Mauritian AAALAC certified breeding centers, were used in this study. All animals were housed in IDMIT facilities (CEA, Fontenay-aux-roses) under BSL-2 containment (Animal facility authorization #D92-032-02, Prefecture des Hauts de Seine, France) and in compliance with European Directive 2010/63/EU, the French regulations and the Standards for Human Care and Use of Laboratory Animals, of the Office for Laboratory Animal Welfare (OLAW, assurance number #A5826-01, US). The protocols were approved by the institutional ethical committee ‘Comité d’Ethique en Expérimentation Animale du Commissariat à l’Energie Atomique et aux Energies Alternatives’ (CEtEA no. 44) under statement number A19_027. The study was authorized by the ‘Research, Innovation and Education Ministry’ under registration number APAFIS #720-201505281237660 v3.

#### NHP Immunizations and sampling

The 25 Cynomolgus macaques were randomly divided into five experimental groups with 5 animals in each. The different vaccine prime-boost regimens for the five groups are illustrated in Figure 3. Animals were sedated with ketamine hydrochloride and medetomidine hydrochloride (10 mg/kg body weight intramuscularly). H107 (20 μg per animal) was administered by the intramuscular (IM) route with adjuvant CAF®01 (625/125μg DDA/TDB), CAF®10b(625/125/50μg DDA/MMG/CpG2006), CAF®09^lo^ (1250/250/62.5μg DDA/MMG/polyI:C), or CAF®09^hi^ (1250/500/125μg DDA/MMG/polyI:C) in 10mM Tris + 4% glycerol, pH 7.0 to a total volume of 0.5mL, at weeks 0 and 4. Control animals received 0.5mL phosphate buffered saline, pH 7.0. All animals were sampled for blood 2 weeks before the first vaccination and at weeks 0, 2, 4, 6, 8, 10, 12 and 30 (Fig. 3).

#### *In vivo* antigen recall in NHPs

During the antigen recall part of the study, six months post the final immunization (see Fig. 3), Purified H107 protein (50 μg per animal in 0.15mL) was administrated by intratracheal route (IT) using endoscope, directly inserted into the trachea until the bifurcation of caudal right lung (accessory lobe and caudal lobe). In parallel, purified H107 and control protein antigen (CTH522; composed of unrelated *Chlamydia trachomatis-specific* antigens) were administrated by intradermal (ID) route consisted of two intradermal injections of 0.1 mL, each containing 20 μg protein antigen per injection the right (H107) or left (control antigen) side of the back of the animal. *In vivo* recall antigens were diluted in 10mM Tris + 4%glycerol, pH 7.0. Pre-medication was performed using alpha-2 agonist atropine sulfate (0.04mg/kg) before anesthesia of the animals to reduce bronchospasm and mucus production during endoscopic exam. Animals were then sedated using ketamine hydrochloride (5 mg/kg, IM) associated with medetomidine hydrochloride (0.05mg/kg IM). After bronchoscopy administration and sampling, animals are injected with Atipamezol hydrochloride (0.25mg/kg) to induce recovery from anesthesia. The animals were sampled for blood at days 0, 3 and 8. On day 8, bronchoalveolar lavage (BAL) was also performed using 50 mL sterile saline, and skin biopsy were performed at injection sites. Blood cell counts, hemoglobin and hematocrit were determined from EDTA-treated blood using an HMX A/L analyzer (Beckman Coulter).

Dermal scoring (skin induration, DTH) was performed at days 0 and 8 post boost. Dermal scoring of the dosing site included observations and graded scoring for erythema, edema, bleeding, scabbing, fissuring and/or ulceration. Skin biopsies were gathered using 8mm^2^ punches and put in PBS after fatty tissue was removed. Briefly, skin biopsies were washed using RPMI 37°C and incubated overnight at 37°C in enzymatic solution (RPMI Glutamax+ 5%FCS+1%ATB+DNase 0.02mg/L) with 4 mg/mL Collagenase D. After incubation, supernatant was collected, aliquoted, and stored at −80°C for antibody ELISA assays. The remaining tissue was dissociated and washed before harvesting cells for stimulation for ICS assays.

#### ELISpot Assays

Fresh PBMCs were used for *ex vivo* stimulation with purified H107 protein or a pool of overlapping 15-mer peptide with (5-10 amino acid overlap) spanning the H107 antigen (*23*) for ELISpot analysis. Briefly, Monkey IFNγ ELISpot PRO kit (Mabtech, Cat #3421M-2APT, Nacka, Sweden) and Monkey IL17A ELISpotPLUS Kit (Mabtech, Cat #3520M-4APW-10) were used following manufacturer’s instructions with 2×10^5^ freshly isolated PBMC added to each well. For IL-17A, H107 protein (5μg/mL) was added in duplicate in the culture medium, and plates incubated ~40hr at +37°C in 5% CO_2_ atmosphere. For IFNγ, in order to reduce non-specific background that was initially observed in pre-immunization screenings, the H107 peptide pool (2μg/mL of each peptide) was used for overnight (~18hr) H107-specific stimulation. Culture medium was used as negative control, and PMA/ionomycin stimulation as a positive control. Plates were then washed 5 times with PBS, followed by addition of biotinylated anti-IFNγ or anti-IL-17A antibodies and a 2h incubation at 37°C. Plates were washed 5 times with PBS, and spots were developed with NBT/BCIP substrate solution. The spots were counted with an Automated Elispot Reader System ELR08IFL (Autoimmun Diagnostika GmbH, Strassberg, Germany).

#### Flow Cytometry

Fresh PBMCs, BAL cells and skin cells were isolated and (1-2 x 10^6^) were resuspended in 150 μl of complete medium containing 0.2 μg of each costimulatory antibody CD28 and CD49d (FastImmune CD28/CD49d, BD). Stimulation was performed in 96 well/plates using H107 protein (5μg/mL), ESAT-6 peptide pool (2μg/mL), H107 peptide pool (1μg/mL) (*23*) or PMA/Ionomycin (as positive control) or culture medium alone (as negative control). Brefeldin A was added to each well at a final concentration of 10 μg/mL and the plate was incubated at 37°C, 5% CO2 overnight. The cells were then washed, stained with a viability dye (dye (LIVE/DEAD fixable Blue dead cell stain kit, ThermoFisher), fixed and permeabilized with the BD Cytofix/Cytoperm reagent. Permeabilized PBMCs and BAL cells were stored at −80°C before the staining procedure. Permeabilized skin samples were directly stained. Staining procedure was performed in a single step following permeabilization using anti-IFNγ (V50), anti-CD4 (BV510), anti-TNFα (BV605), anti-IL-13 (BV711), anti-CD154 (FITC), anti-IL-2 (PerCP/Cy5.5), anti-CD8 (PE-Vio770), anti-IL-17A (Alexafluor 700), CD3 (APC-Cy7). After 30 minutes of incubation at 4°C in the dark, cells will be washed in BD Perm/Wash buffer. Cells were acquired with an LSR II (BD) after the staining procedure. FlowJo software v10 (TreeStar) was used for sample analysis. PCA analysis was performed using GraphPad Prism v9 software on Boolean gating data normalized to percentage of the total cytokine positive population per individual animal.

#### PET-CT

Chest PET-CT was performed at baseline prior to antigen recall (D0) and on days 3 (D3) and 7 (D7) post recall. Animals were fasted for at least 8 h before each imaging session. All imaging acquisition was performed using the Digital Photon Counting (DPC) PET-CT system (Vereos-Ingenuity, Philips) implemented in a BSL-3 laboratory. These sessions were always performed in the same experimental conditions (acquisition time and animal order) to limit [18F]-FDG-PET experimental bias. Animals were first anesthetized with ketamine hydrochloride (5 mg/kg, IM) associated with medetomidine hydrochloride (0.05mg/kg IM), intubated, and then maintained under 0.5-1.5% isoflurane and placed in a supine position on a warming blanket (Bear Hugger, 3M) on the machine bed with monitoring of the cardiac rate, oxygen saturation, and body temperature.

Computed Tomography (CT) was performed 5 minutes prior to PET acquisition for attenuation correction and anatomical localization. The CT detector collimation used was 64 × 0.6 mm, the tube voltage was 120 kV, and the intensity was approximately 150 mAs. Chest-CT images were reconstructed with a slice thickness of 1.25 mm and an interval of 0.63 mm. A whole-body PET scan (3 bed positions, 3 min/bed position) was performed approximately 45 min post-injection of 3.5 ± 0.4 MBq kg-1 of [18F]-FDG via the saphenous vein. PET images were reconstructed onto a 256 x 256 matrix using OSEM (3 iterations, 15 subsets). After PET acquisition, animals were resuscitate using atipamezole hydrochloride (0.25mg/kg).

PET images were analyzed using 3DSlicer software (open-source tool). For segmentation, various regions of interest (entire lung and separated lung lobes) were semi-automatically contoured according to anatomical information from CT. A 3D volume of interest (VOI) was interpolated from several ROIs in different image slices to cover each lung lobe excluding background signal (heart and liver). For quantification, [18F]-FDG accumulation in the VOIs was given as a standardized uptake value (SUVmean, SUVmax).

#### H107-specific Antibody ELISAs in NHPs

An indirect quantitative ELISA was developed to measure the content of anti-H107 IgG and IgA antibodies in NHP sera and BAL samples. Maxisorb Plates (Nunc) were coated overnight with purified H107 antigen (0.1μg/ml) at 4°C, and then >1.5 hrs blocked with 2% BSA at room temperature. Sera were diluted 1:200 (1:20 from control animals) and thereafter seven 2-fold serial dilutions made. Undiluted BAL and thereafter seven 3-fold serial dilutions were made. Sample dilutions were added to the plates and incubated overnight (~16hrs) at 4°C. HRP-conjugate anti-IgG (goat anti-Monkey IgG, Fitzgerald Cat# 43R-IG020HRP, at 1:5000 dilution) or IgA (polyclonal rabbit anti-human IgA/HRP, Dako, P0216, at 1:2000 dilution) was added for 1 hr at room temperature and the enzyme reaction was developed with TMB plus (Kementec) for 15 minutes and stopped with 0.2 M H_2_SO_4_ solution. Plates were read at 450 nm with 620 nm background correction using an ELISA reader (Tecan Sunrise). We confirmed the specificity of our IgA ELISA and non-cross reactivity toward IgG using purified human IgG and IgA (Sigma-Aldrich).

A reference serum pool was established by combining sera from all immunized NHP from study week eight and arbitrarily assigned a value of 10,000 ELISA-Units (AU). Two-fold serial dilutions of reference sample was run on each plate. For each sample, titers were calculated from five parameter logistic curves based on 2-fold dilution curves using the package ‘drc’ in R. Sample titer values were based on dilutions from the linear portion of the response curve for IgG and IgA assays. Area under the curve (AUC) calculation was performed in GraphPad Prism v9 after baseline correction.

#### Luciferase Assay

THP-1-Dual cells (InVivoGen) were grown under selection media at 37°C according to manufacturer’s instruction. 10^5^ cells were seeded per well of 96-well flat bottom plate in triplicate and differentiated for 48hrs with PMA (40ng/ml). PMA was removed and CAF® adjuvants or varying concentrations added to the cells for overnight incubation at 37°C. Supernatants were collected and secreted Luciferase activity assayed using QUANTI-Luc™ kit (InVivoGen) as per manufacturer instructions. The CyQUANT-LDH-Cytotoxicity assay (Invitrogen) was also perform on supernatants according to manufacturer instructions, confirming that cell viability was not increased compared to untreated cells. Assays were read using a SpectraMax iD3 (Molecular Devices).

#### Statistics and Illustrations

All statistical tests and PCA analyses were calculated using the statistical Graphpad Prism software v9. Figure illustrations were created using Biorender.com.

## List of Supplementary Materials

Fig. S1 to S6.

## Acknowledgments

We thank Peter Andersen for his discussions and contribution to study design. We thank Rune Fledelius Jensen for preparation of CAF® adjuvants, Vivi Andersen for H107 protein production, and Sara Knudsen for technical support of antibody data analysis. We thank the entire Animal facility staff at SSI for their technical expertise, skills and efforts. We thank the staff of the animal facility of IDMIT, particularly J. Lemaitre, B. Delache, M. Pottier, JM. Robert and Francis Relouzat, the staff of FlowCytech laboratory, particularly, M. Gomez-Pacheco, the Immunology and Infectiology teams particularly, L. Bossevot, M.Galpin-Lebreau, M. Leonec and L.Moenne-Loccoz, and R. Marlin, C. Bouillier and I. Mangeot from IDMIT for their technical expertise, skills and efforts. The Infectious Disease Models and Innovative Therapies research infrastructure (IDMIT) is supported by the “Programmes Investissements d’Avenir” (PIA), managed by the ANR under references ANR-11-INBS-0008 and ANR-10-EQPX-02-01. We thank the Fondation Dormeur Vaduz for the donation of instruments relevant to this project.

## Author contributions

Conceptualization: JSW, RM, DC, TL, GKP

Methodology: JSW, VC, AG, TN, NB, MLO, IR, RLG, GKP, RM

Resources: DC, IR, GKW

Investigation: JSW, VC, TN, NK, AG, SL, EB, CJ, WS, JM, MLO, AS, TL

Formal Analysis: JSW, VC, TN, AG, WS, GKW, GKP, RM

Visualization: JSW, VC, TN, AS, GKW, GKP, RM

Funding acquisition: RM, FF

Project administration: JSW, VC, RLG, RM

Supervision: FF, TL, RLG, GKP, RM

Writing – original draft: JSW, RM

Writing – review & editing: VC, DC, TN, GKW, FF, GKP

