## Supplemental Figures for "A novel adjuvant formulation induces robust Th1/Th17 memory and mucosal recall responses in Non-Human Primates"

### Supplemental Materials

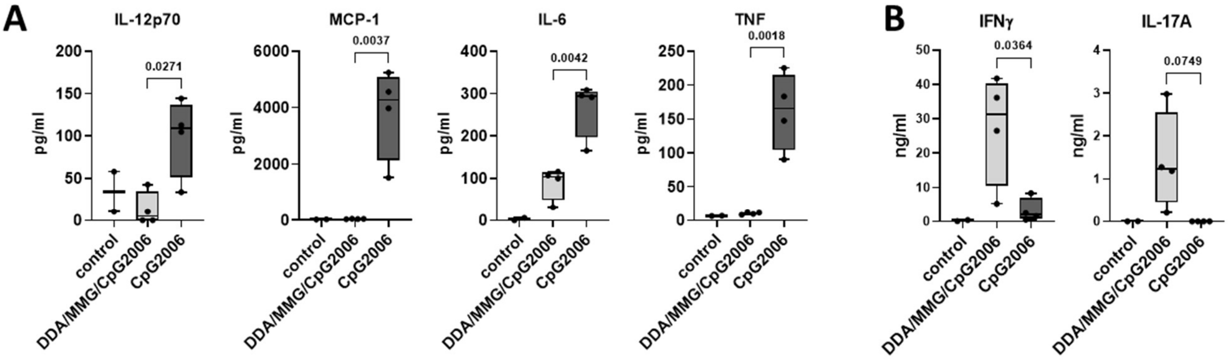


**Figure S1. Incorporation of high dose CpG into DDA/MMG liposomes reduces its toxicity and promotes a Th17 response.**

CB6F1 mice immunized 2x SC with H56 protein antigen adjuvanted with 50µg CpG2006 or 50µg CpG formulated into DDA/MMG liposomes (250/50 µg) or left unimmunized (control). **(A)** Serum levels of inflammatory cytokines and chemokines TNF, IL-6, IL-12-p70, and MCP-1 measured 2 days after the first immunization. **(B)** Splenocytes taken 3 weeks after the final immunization and assessed by ELISA for secretion of IFNγ (left) and IL-17A (right) after *ex vivo* stimulation with H56 protein. Symbols represent individual mice. Bars represent median, IQR, min. and max values. P values, one-way ANOVA with Tukey's posttest.

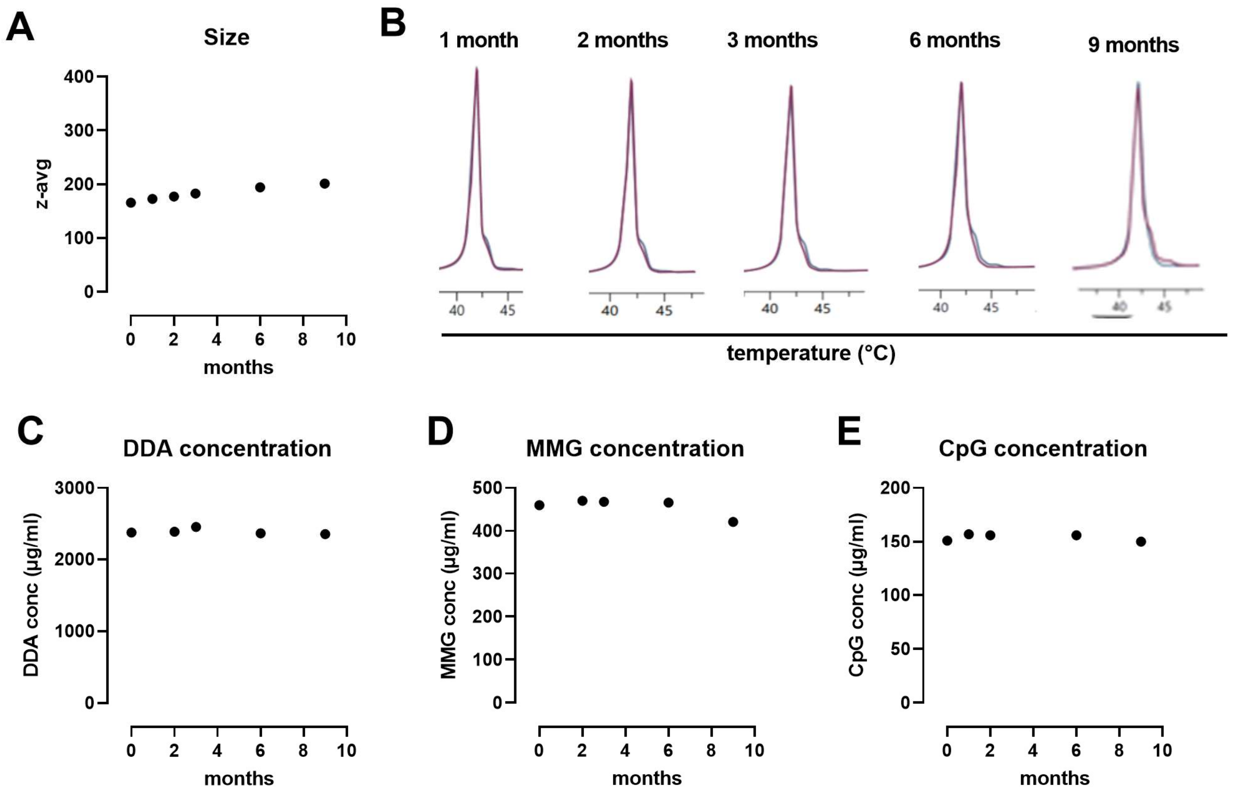


**Figure S2. Long-term stability of DDA/MMG/CpG liposomes**

Long-term stability of CAF®10b liposomes was assessed for 9 months. **(A)** Size of the liposomes as measured by dynamic light scattering. **(B)** thermotropic phase behavior of the liposomes in suspension. Content of **(C)** DDA, **(D)** MMG, and **(E)** CpG2006 when stored at 4 degrees Celsius. (A,C-E) Symbols indicate mean of three measurements.

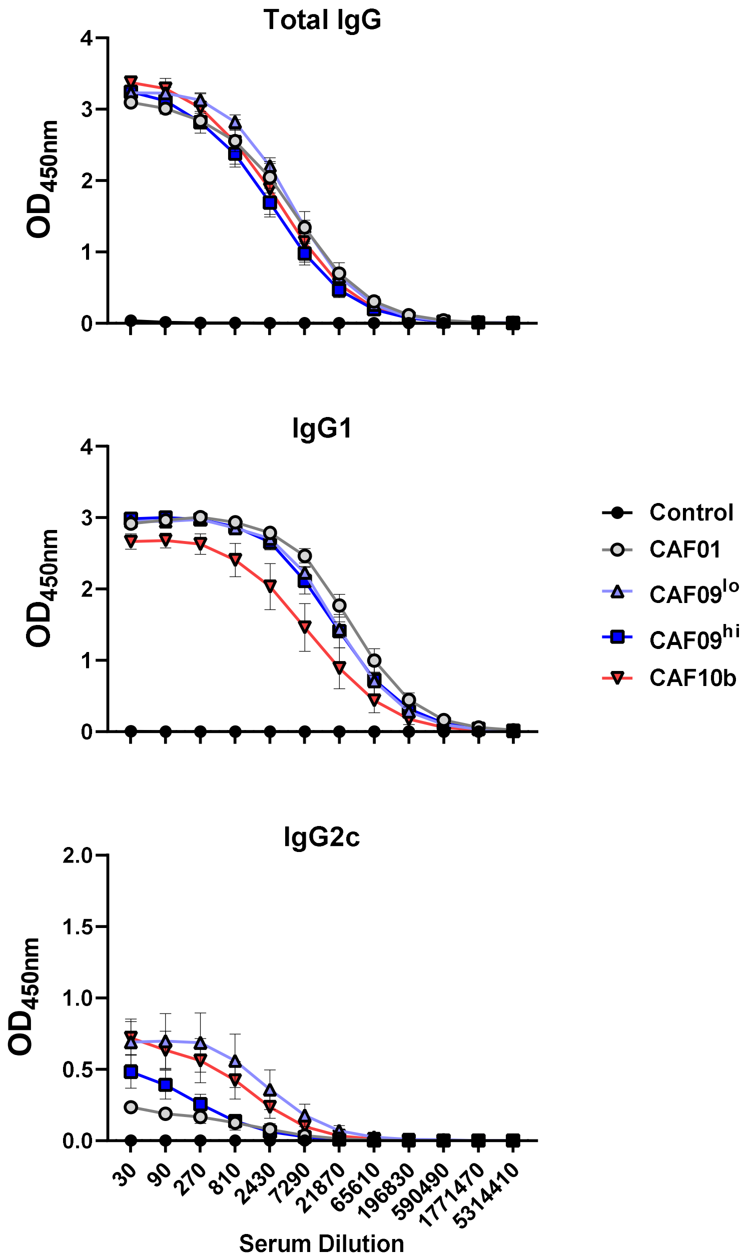


**Figure S3. Vaccine specific serum antibody responses in mice**

H107-specific serum antibodies one day after final immunization of CB6F1 mice. Total IgG (top), IgG1 (middle), IgG2c (bottom) measured by ELISA and $OD_{450}$ shown for the indicated serum dilutions. Symbols indicate mean ± SD.

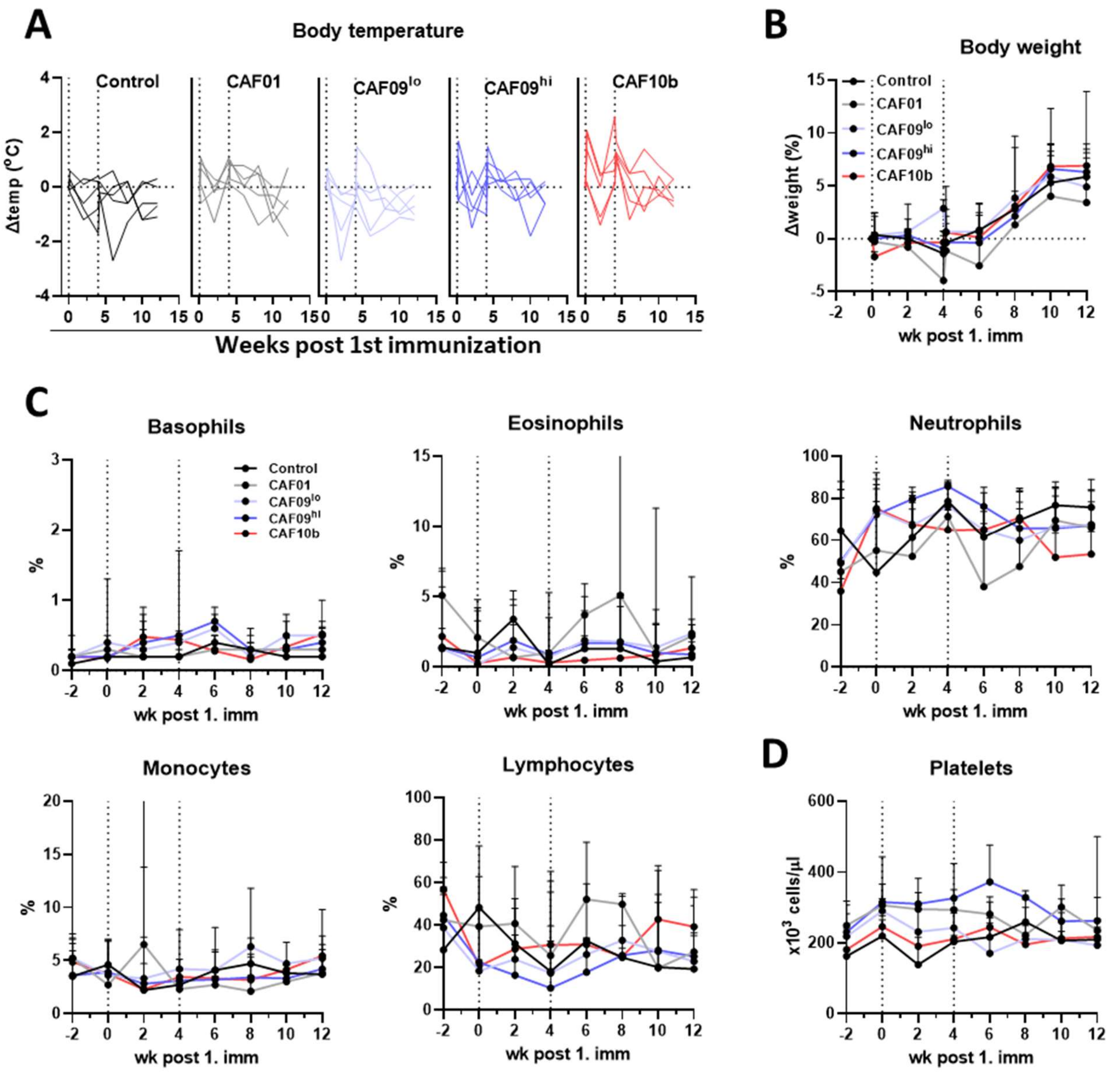


**Figure S4. Vitals and CBC analysis of NHPs after CAF® immunizations**

NHPs were monitored for changes in vital signs and blood cellular composition after immunizations. **(A)** body (rectal) temperature, shown as change from baseline (wk 0). Lines indicate individual animals. **(B)** Body weight depicted as percent change from baseline. **(C)** Percent of total white blood cells depicted over time for each cell type indicated. **(D)** Platelet count over time indicated. (B-D) Symbols indicate median±range per group.

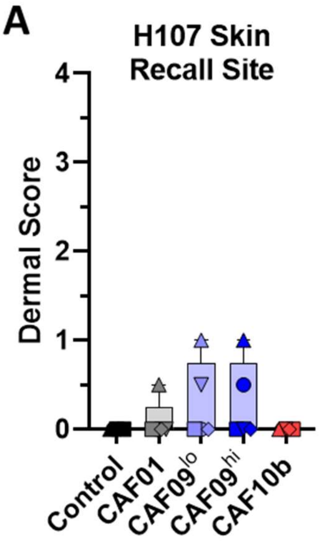


| Score | Grade | Edema | Erythema |
|---|---|---|---|
| 0 | None | No swelling | Normal color |
| 1 | Minimal | Slight swelling, indistinct border | Light pink; indistinct |
| 2 | Mild | Defined swelling, distinct border | Bright pink; distinct |
| 3 | Moderate | Defined swelling, raised border (<1mm) | Bright red; distinct |
| 4 | Severe | Defined swelling, raised border (≥1mm) | Dark red; pronounced |

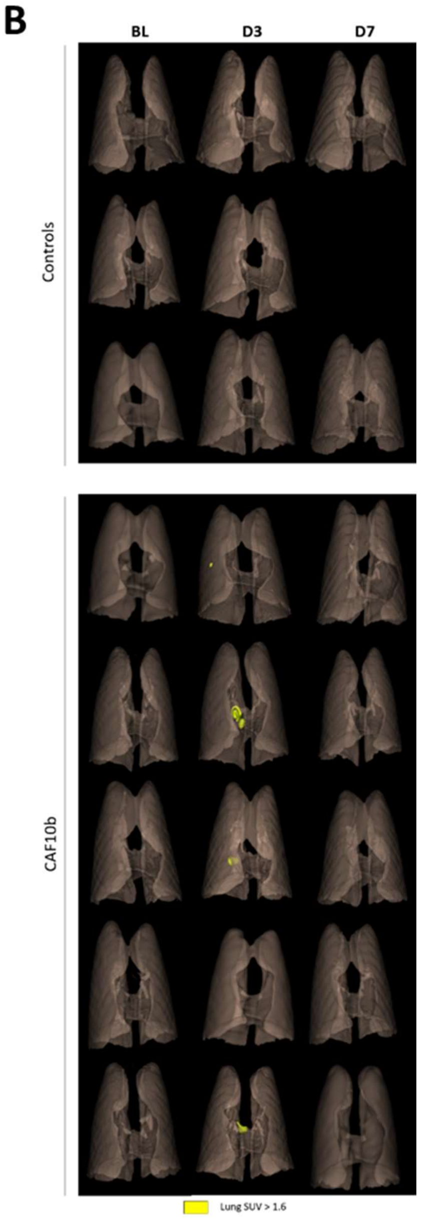


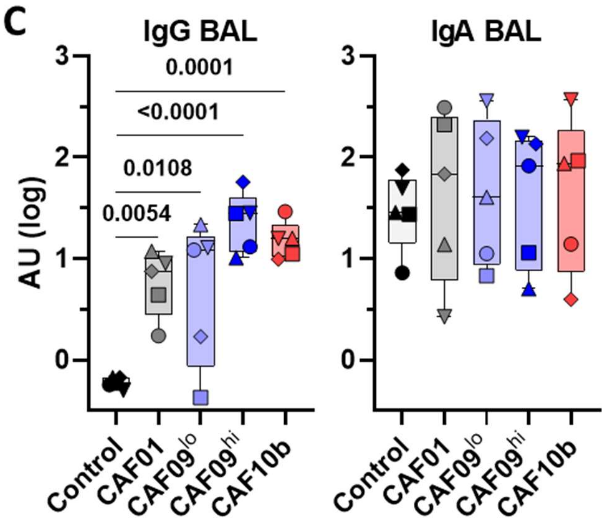


**Figure S5. *In vivo* Antigen recall analyses in NHPs**

NHPs were analyzed after ID+IT H107 antigen recall. **(A)** Local Skin reactions at H107-ID site day 8 post antigen recall. Symbols indicate the average score of the two H107 antigen sites on each animal. **(B)** PET-CT performed on selected NHPs (n=3 Control and n=5 H107/CAF®10b animals) before (baseline, BL) and 3 and 7 days after instillation of H107 protein into the right lung. The SUV maximum fold change for each animal is shown. **(C)** H107 specific IgG (left) and IgA (right) in the BAL collected from the right lung, 8 days after instillation of H107 protein. AU, arbitrary units. Bars represent median ± IQR, min. and max. Symbols, individual animals. P values from ANOVA with Dunnett's posttest of log-transformed data.

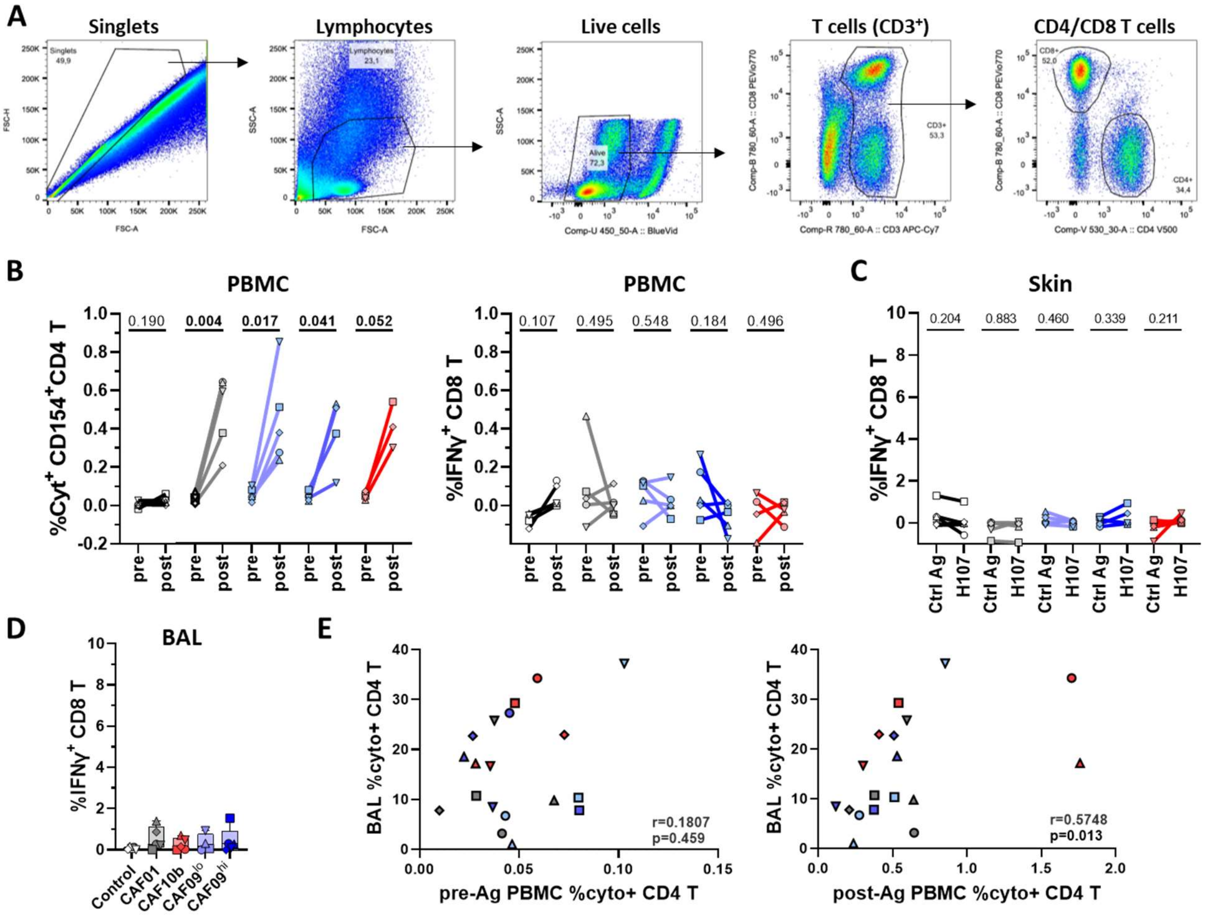


**Figure S6. Cellular analyses of in vivo antigen recall in NHPs**

Cells isolated from blood, skin, and BAL were analyzed by intracellular cytokine staining (ICS). **(A)** Gating strategy of CD4 and CD8 T cells for analysis of cytokine production by ICS from PBMC, BAL, and skin biopsy cells. Example shown is BAL after antigen recall. **(B)** The percentage of cytokine-producing CD4 T cells (left) and CD8 T cells (right) from PBMC samples collected pre- and post-*in vivo* antigen recall and assessed IFNγ, TNFα, IL-2, IL-17, and/or IL-13 after *ex vivo* stimulation with H107 protein with subtraction of non-stimulated background (-NS) per animal. **(C)** The percentage of IFNγ-producing CD8 T cells isolated from skin biopsies collected from control antigen (Ctrl Ag) and H107 ID injection sites after *ex vivo* stimulation with H107 protein (-NS). (B,C) Symbols and lines, individual animals. P values from paired T tests. **(D)** The percentage of H107-specific cytokine-expressing BAL CD8 T cells as determined by ICS (-NS). Bars represent median ± IQR, min. and max. Symbols, individual animals. Kruskal-Wallis test p=0.6247. (E) Scatter plots of H107-specific CD4 T cell responses (%cyto+ by ICS) in PBMCs pre- and post-*in vivo* recall versus BAL responses in H107/CAF® immunized NHP. Symbols indicate individual animals; CAF®01(grey), CAF®09$^{lo}$(light blue), CAF®09$^{hi}$(dark blue), CAF®10b(red). Spearman correlation coefficients (r) and p values are shown.
